# Machine learning unveils latent architecture of superiority illusion that predicts visual illusion perception and metacognitive performance

**DOI:** 10.1101/2022.10.24.513462

**Authors:** Daisuke Matsuyoshi, Ayako Isato, Makiko Yamada

## Abstract

Humans are typically inept at evaluating their abilities and predispositions, while often disregarding such lack of metacognitive insight into their capacities and even augmenting (albeit illusorily) self-evaluation such that they should have more desirable traits than an average peer. This superiority illusion helps maintain a healthy mental state. However, the scope and range of its influence on broader human behavior, especially perceptual tasks, remain elusive. As belief shapes the way people perceive and recognize, the illusory self-superiority belief potentially regulates our perceptual and metacognitive performance. In this study, we used hierarchical Bayesian estimation and machine learning of signal detection theoretic measures to understand how superiority illusion influences visual perception and metacognition for Ponzo illusion. Our results demonstrated that superiority illusion correlated with visual illusion magnitude and metacognitive performance. Next, we used machine learning with a relaxed elastic net and unveiled the latent architecture that underlies the correlations. We revealed that the “extraversion” superiority dimension tapped into visual illusion magnitude and metacognitive ability. In contrast, the “honesty-humility” and “neuroticism” dimensions were only predictive of visual illusion magnitude and metacognitive ability, respectively. These results suggest common and distinct influences of superiority features on perceptual sensitivity and metacognition. Our findings contribute to the accumulating body of evidence indicating that the superiority illusion leverage is far-reaching, even to visual perception.

**Significance Statements:** People have a cognitive bias to overestimate their abilities above the mean (superiority illusion) and thereby help maintain a healthy mental state. In this work, we show that the influences of superiority illusion are more extensive than previously thought. We find that superiority illusion correlated with visual illusion magnitude and metacognitive performance. Furthermore, using hierarchical Bayesian estimation and machine learning, we unveil the latent architecture (i.e., overlapping yet dissociable superiority features) that predicts visual illusion magnitude and metacognitive performance. These findings suggest that superiority illusion is a cardinal cognitive bias that involves a vast assortment of behavior as an illusion is an efficient and adaptive tool for humans to somehow thrive in a world of ambiguity.

## Introduction

Contrary to our naïve belief, humans often do not have accurate insight into themselves. The metacognitive capacity to assess self-made decisions or personal abilities varies substantially across individuals, typically ranging 30–100% of the information theoretically available to an individual (1, 2). People dismiss their lack of metacognitive insight and even increase self-evaluation such that they should have more desirable traits than an average peer (3-5). At first glance, this superiority illusion (SI) appears as a metacognitive ability defect. However, evidence suggests that SI helps maintain a healthy mental state (3, 4, 6, 7), self-esteem (8, 9), and life satisfaction (9, 10) except for overly optimistic self-evaluations (11-14). Therefore, rather than a defect, SI is likely to be a self-serving cognitive bias with a myriad of psychological benefits.

Numerous studies have shown that SI occurs in various domains (15-17) with cross-cultural robustness (18-21), indicating its universal and fundamental contributions to human behavior. However, the scope and range of how SI influences perception remains unclear. As cognitive style underlies the way people perceive, think, solve problems, learn, and relate to others (22), SI also likely exerts its heuristics, even over perception, by biasing cognition and decision-making toward illusory ones. For example, field-dependence/independence is among the best-known cognitive styles (23). Field-dependent people are known to be inept at absolute size estimation (24) and various visuospatial tasks (25) and are susceptible to the Ponzo illusion (26). Although Zhang claimed that the field-dependence/independence construct represents a perceptual ability rather than a cognitive style (27), later studies demonstrated that cognitive styles represent behavioral heuristics that govern across multiple levels of information processing, from perceptual ability and metacognition to personality traits and social skills (28-31).

In this study, we used hierarchical Bayesian estimation and machine learning of signal detection theoretic (SDT) measures to understand how SI influences visual illusion. As retinal images are inherently ambiguous (e.g., a distant large or a closer small object could invoke the same retinal projection), human vision resolves ambiguities by biasing neural activities based on knowledge or beliefs (32, 33). We hypothesized that visual illusion is a powerful window into how we incorporate various sources and create best-bet predictive hypotheses of objects and situations for optimal, adaptive behavior while handling environmental uncertainties. We chose the Ponzo illusion as a visual stimulus (34) since it must be mediated by feedback projections from higher areas and is prone to top–down control (35, 36). Compared to other illusions established only by lateral connections in the primary visual cortex (37-39), these characteristics of the Ponzo illusion are desirable for our study investigating the effects of top–down, illusory cognitive bias. To examine visual illusion perception and its metacognition in the SDT framework, unlike a typical experiment using a method of adjustment or constant stimuli, we asked participants whether the two stimuli were the same or different (same/different task) and to rate their metacognitive confidence about the perceptual decision (confidence rating task) (Fig. 1). Although a perceptually-demanding task (i.e., visual illusion) often prevents us from estimating reliable metacognitive ability (1, 40), hierarchical Bayesian estimation allows for accurately estimating metacognitive measures even when expecting low perceptual sensitivity (41).

**Figure 1.**
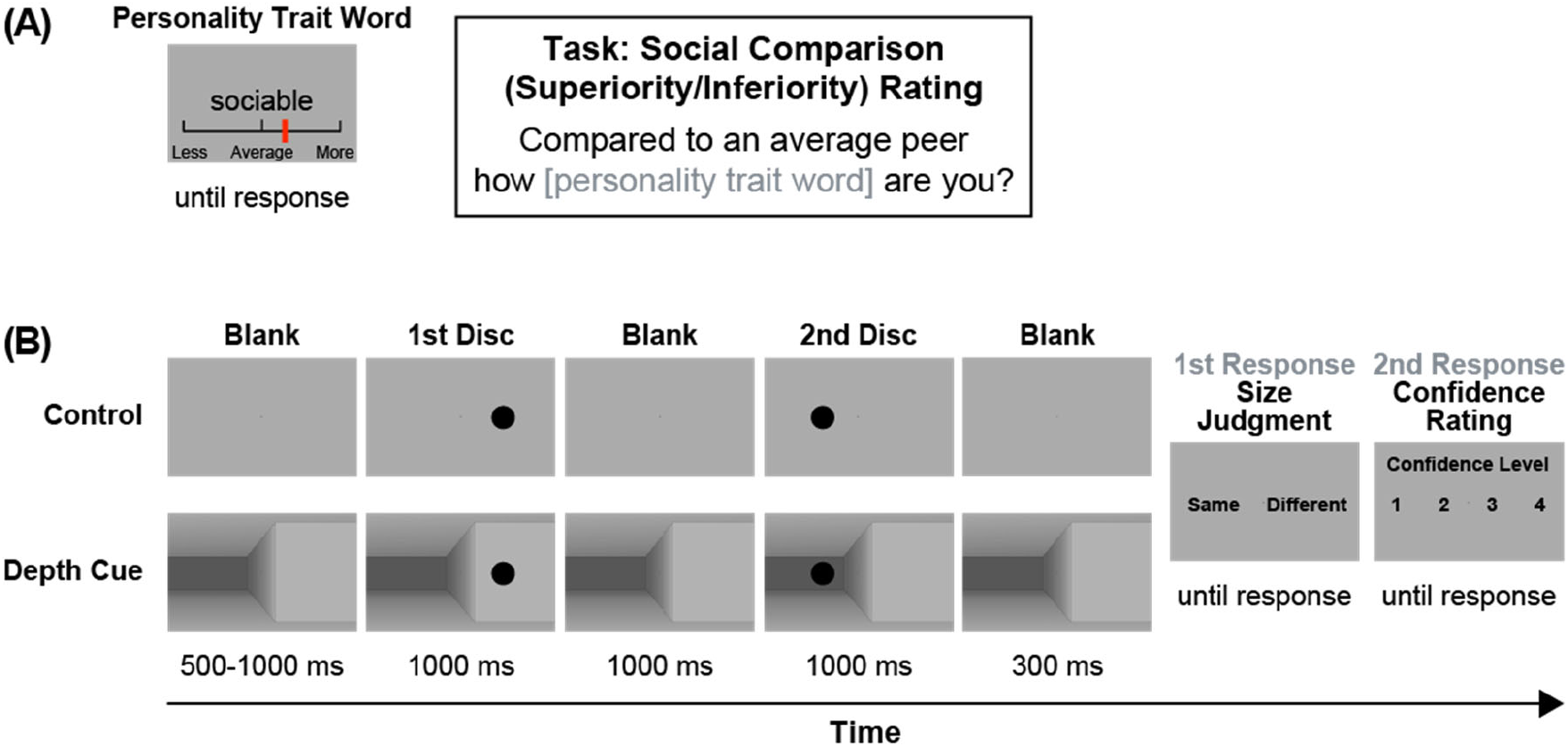
Experimental paradigm. (A) Schematic presentation of the superiority rating task. Participants indicated how personality trait words described them compared to an average peer using a sliding scale. (B) Schematic presentation of the Ponzo illusion task. Participants were required to indicate whether the two discs were of the same size (1st response), then rate their confidence (2nd response).

Moreover, we combined machine learning and principal component analysis (PCA) to create prediction models for visual illusion magnitude and metacognitive performance from SI rating data. This combined approach allowed us to uncover the latent architecture underlying the models by examining the product of machine-learning feature importance and SI PCA loadings. The current machine learning approach focuses on effectively extracting hidden information in the data rather than simply creating prediction models. The approach presented in this study enables us to gain an in-depth understanding of behavioral findings by unveiling how latent SI features influence visual perception and metacognition.

## Results

### Superiority rating

We asked participants to rate their superiority/inferiority compared to an average peer. The mean superiority rating score was 0.0815, significantly greater than zero (*t*_*36*_ = 2.6334, *p* = 0.0124, Cohen’s *d* = 0.4329 [95% CI: 0.0995, 0.7663]), confirming the superiority bias of the participants toward their own abilities or traits (Fig. 2A).

**Figure 2.**
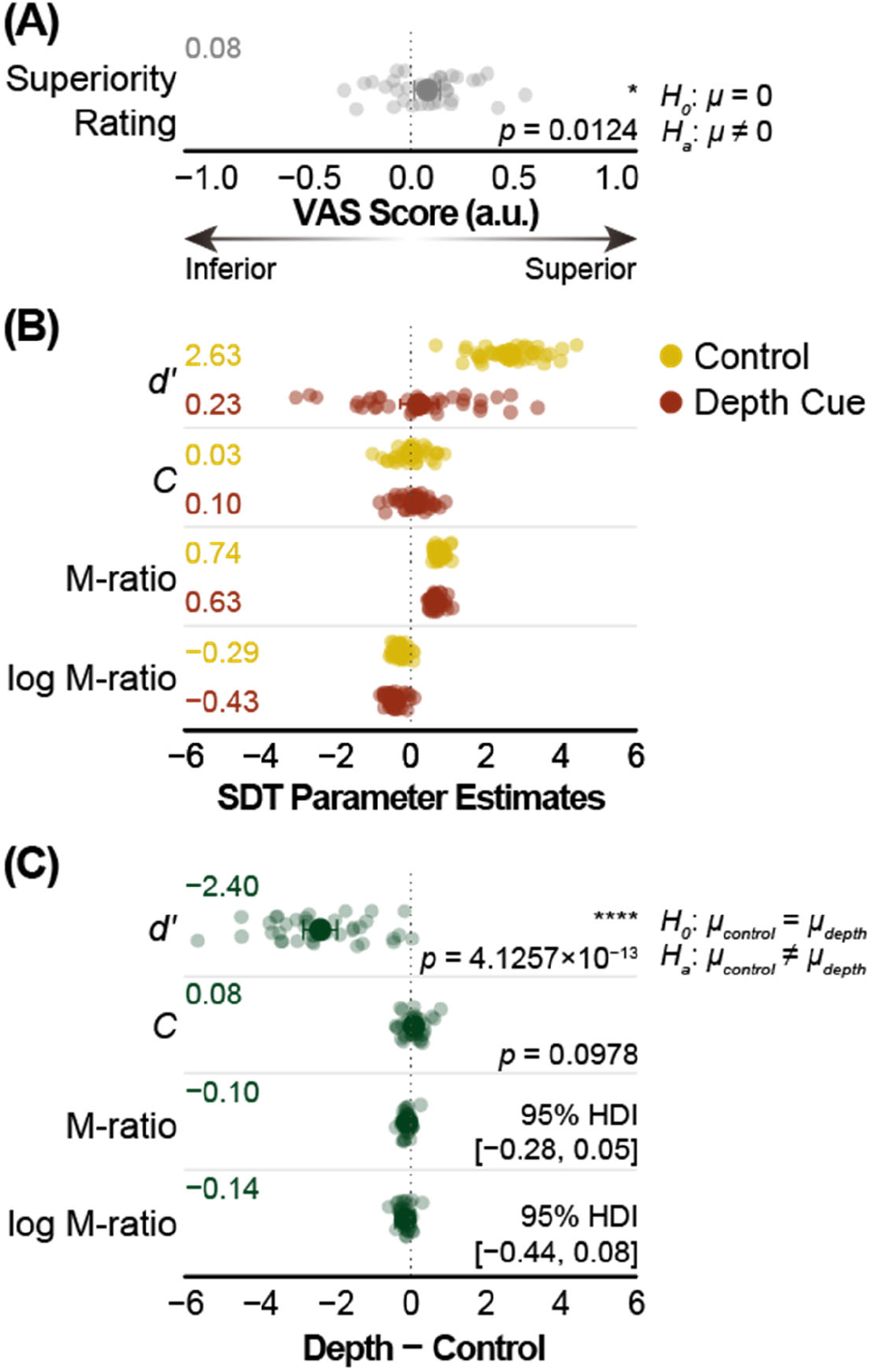
Behavioral results. (A) Superiority rating score. (B) Signal detection theoretic (SDT) parameter estimates for the Ponzo illusion task. (C) SDT parameter estimate differences (between depth cue and control conditions) for the Ponzo illusion task. In A-C, transparent dots represent individual data points (superiority rating and type 1 SDT) or individual estimates obtained from a single-subject Bayesian model fit (type 2 SDT). Larger non-transparent dots and corresponding leftmost values represent the mean values across participants (superiority rating and type 1) or the group-level hierarchical Bayesian maximum a posteriori (MAP) probability estimates (type 2). Error bars represent 95% confidence intervals of the mean (superiority rating and type 1) or 95% highest-density intervals (HDI) of posterior distributions (type 2). The rightmost values indicate statistical test values. Asterisks represent statistical significance (* *p* < 0.05, **** *p* < 0.0001). M-ratio = meta-*d’* / *d’*. log M-ratio = log(meta-*d’* / *d’*). VAS, visual analog scale. a.u., arbitrary unit.

### Ponzo illusion

Figure 2B presents the SDT parameter estimates for the Ponzo illusion. In the case of type 1 estimates, one-sample t-tests indicated significantly larger than zero *d’* values under control conditions (*t*_*36*_ = 18.7067, *p* = 3.9530 × 10^−20^, Cohen’s *d* = 3.0754 [95% CI: 2.7419, 3.4088]), while *d’* was comparable to zero under depth cue conditions (*t*_*36*_ = 0.9052, *p* = 0.3714, Cohen’s *d* = 0.1488 [95% CI: −0.1846, 0.4822]). In addition, criterion *C* was comparable to zero under both control (*t*_*36*_ = 0.360382, *p* = 0.7207, Cohen’s *d* = 0.0592 [95% CI: −0.2742, 0.3927]) and depth cue (*t*_*36*_ = 1.5603, *p* = 0.1274, Cohen’s *d* = 0.2565 [95% CI: −0.0769, 0.5899]) conditions.

In the case of type 2 estimates, we performed a hierarchical Bayesian estimation of metacognitive parameters from confidence ratings (41). The group-level hierarchical Bayesian maximum a posteriori (MAP) probability M-ratio estimates were 0.7441 and 0.6276 (control and depth cue conditions, respectively). They were smaller than one under both control (95% HDI: 0.6498, 0.8416) and depth cue (95% HDI: 0.6498, 0.8416) conditions. Log M-ratio MAP estimates were −0.2921 and −0.4325 (control and depth cue conditions, respectively). They were smaller than zero under both control (95% HDI: −0.4253, −0.1669) and depth cue (95% HDI: −0.7357, −0.2489) conditions, indicating that metacognitive monitoring is not optimal for either task.

Figure 2C shows the between-condition differences (depth cue − control) in the SDT parameter estimates for the Ponzo illusion. Under depth cue conditions, smaller *d’* values could be obtained than control conditions (*t*_*36*_ = 11.0415, *p* = 4.1257 × 10^−13^, Cohen’s *d* = −1.8152 [95% CI: −2.1486, −1.4818]), confirming that strong Ponzo illusion was induced by depth cues. Criterion *C* was comparable between the depth cue and control conditions (*t*_*36*_ = 1.6997, *p* = 0.0978, Cohen’s *d* = 0.2794 [95% CI: −0.0540, 0.6128]). We found a non-meaningful between-condition differences for the M-ratio (MAP = −0.1049, 95% HDI: −0.2788, 0.0517) or log M-ratio (MAP = −0.1372, 95% HDI: −0.4371, 0.0795), indicating the metacognitive performance of the participants was comparable between the conditions.

### Correlations between superiority rating, perceptual sensitivity, and metacognitive efficiency

Figure 3 shows correlations between superiority rating, perceptual sensitivity (*d’*), and metacognitive performance (log M-ratio) scores. Both *d’* (*rho* = −0.4009, *p* = 0.0139) and log M-ratio (*rho* = −0.4594, *p* = 0.0042) significantly correlated with superiority rating scores (Fig. 3A), while no significant correlation could be found between the *d’* value and log M-ratio (Fig. 3B, *rho* = 0.2245, *p* = 0.1810). These results remained constant even when controlling for each other. We detected significant partial correlations between *d’* and superiority rating scores (*rho*_*p*_ = −0.3440, *p* = 0.0399) and between log M-ratio and superiority rating scores (*rho*_*p*_ = −0.4137, *p* = 0.0121) while controlling for the log M-ratio and *d’*, respectively. In addition, we found a nonsignificant correlation between criterion *C* and the superiority rating scores (*rho* = 0.1718, *p* = 0.3093).

**Figure 3.**
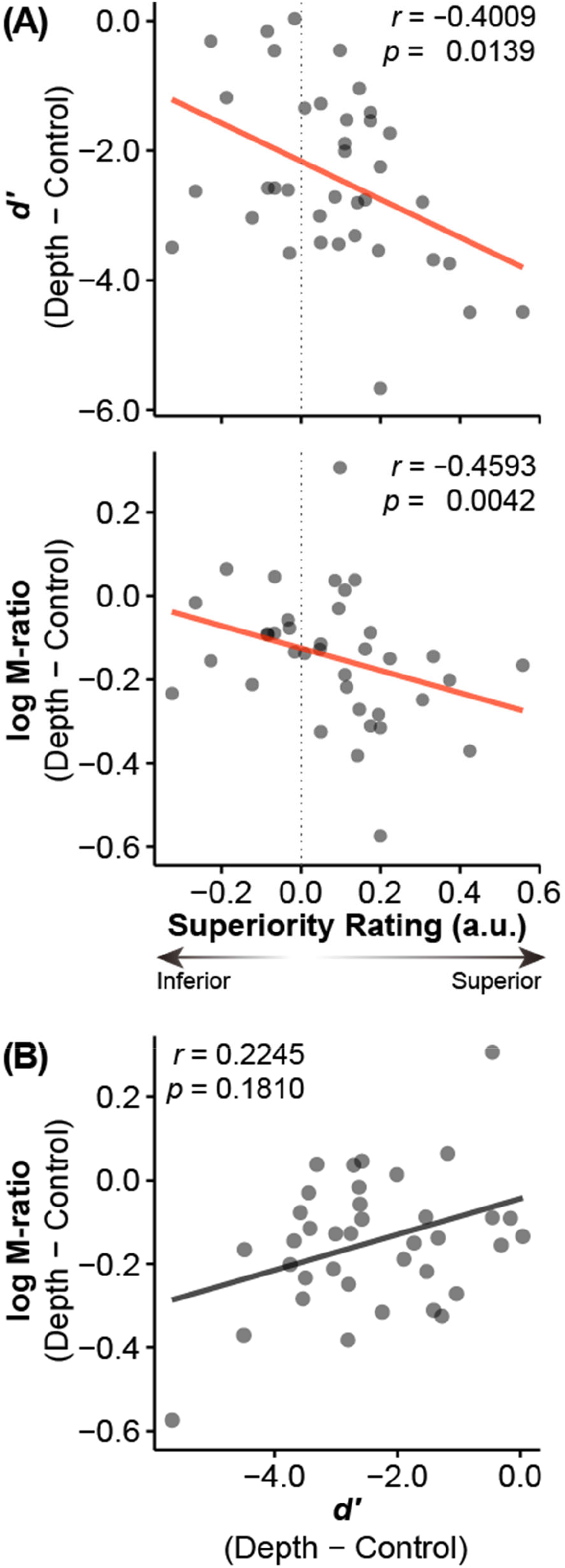
Correlations between superiority rating, perceptual sensitivity (*d’*), and metacognitive efficiency scores (log M-ratio). (A) Both the *d’* value and log M-ratio exhibited significant correlations with superiority rating scores. (B) No significant correlation between *d’* and log M-ratio. Transparent dots represent individual data points. Transparent lines represent linear regression fit using ordinary least squares. a.u., arbitrary unit.

### Machine learning model

The relaxed elastic net regression with leave-one-sample-out cross validation (LOOCV) revealed that different sets of superiority rating items predicted each SDT parameter estimate (Table1). Both the *d’* and log M-ratio models achieved good accuracy (*rho* = 0.7205, *p* = 1.4369 × 10^−6^, *R*^*2*^ = 0.5341, *root-mean square error* [*RMSE*] = 0.6735; *rho* = 0.6700, *p* = 1.0028 × 10^−5^, *R*^*2*^ = 0.3909, *RMSE* = 0.7708) and consisted of seven and six superiority rating items (Fig. 4 top and bottom row), respectively. No overlap could be observed between the two machine learning model items, indicating that the *d’* and log M-ratio parameter estimates were independently correlated (at least in part) with superiority ratings.

**Figure 4.**
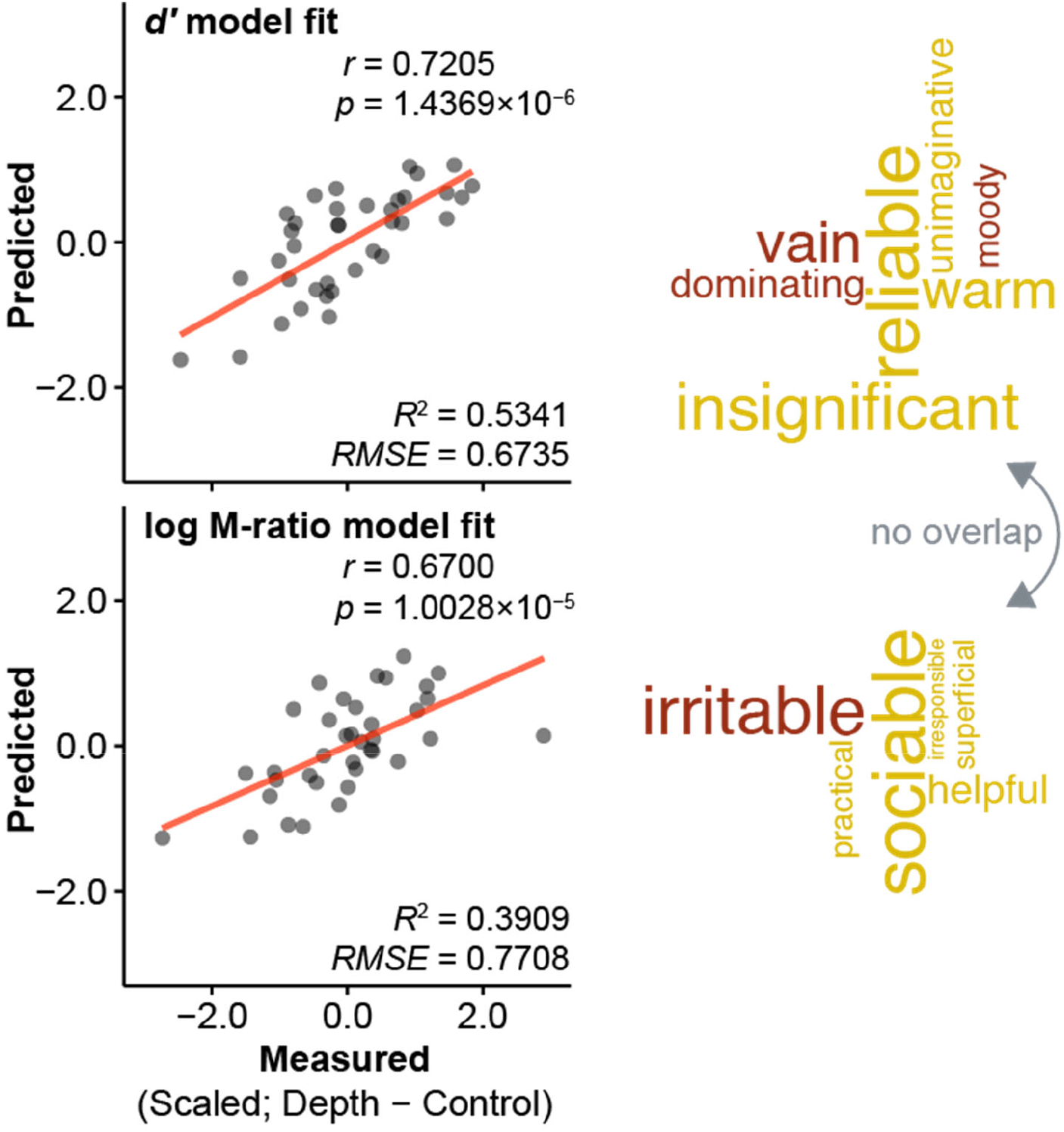
Machine learning prediction of perceptual sensitivity (*d’*) and metacognitive efficiency (log M-ratio) from superiority rating scores. Relaxed elastic net regression with eave-one-sample-out cross-validation created prediction models for *d’* (top row) and log M-ratio (bottom row) from superiority rating scores. Although the two models displayed similar prediction accuracy (left column), they consisted of different superiority rating items (right column). For more information, refer to Table 1. Transparent dots represent individual data points. Transparent lines represent linear regression fit using ordinary least squares. The word size was scaled relative to (absolute value of) machine learning feature importance in the word cloud plot. Red and yellow words denote positive and negative feature importance, respectively (Table 1). *R*^*2*^, r-squared. *RMSE*, root-mean square error.

### Latent architecture underlying machine learning model items

Given that the machine learning models selected different items for each model, it is possible that *d’* and log M-ratio *completely* independently correlated with superiority ratings. However, an identical latent component might underlie the correlations even if the two models contained different items. To examine this possibility, we performed a PCA on 52-item superiority ratings, then assessed the relative contribution of each PC to each model.

**Table 1.** Feature importance in machine learning models for predicting perceptual sensitivity (*d’*) and metacognitive efficiency (log M-ratio) based on superiority rating scores.

| Item |  | Feature importance |
| --- | --- | --- |
| *d'* |  |  |
| #29 | vain | 0.2026 |
| #21 | dominating | 0.1276 |
| #05 | moody | 0.1073 |
| #07 | unimaginative | -0.1185 |
| #08 | warm | -0.1942 |
| #13 | insignificant | -0.2372 |
| #35 | reliable | -0.2656 |
| *log M-ratio* |  |  |
| #19 | irritable | 0.2990 |
| #48 | irresponsible | -0.0615 |
| #33 | superficial | -0.1008 |
| #32 | practical | -0.1159 |
| #50 | helpful | -0.1586 |
| #45 | sociable | -0.3227 |

Note that response and predictor variables included in the machine learning models were standardized for each variable, so the model feature importance should be interpreted accordingly. For prediction performance, see Figure 4.

The PCA with parallel analysis (42) revealed three significant PCs underlying the 52-item superiority ratings (Table 2). PC1 consisted of items such as “sociable” and “reliable,” so we labeled this PC as the “*extraversion*” component. PC2 consisted of items such as “persistent” and “honest,” this PC might thus, reflect the “*honesty-humility*” component. PC3 consisted of items such as “sentimental” and “irritable,” we thus, regarded this PC as the “*neuroticism*” component.

**Table 2.**
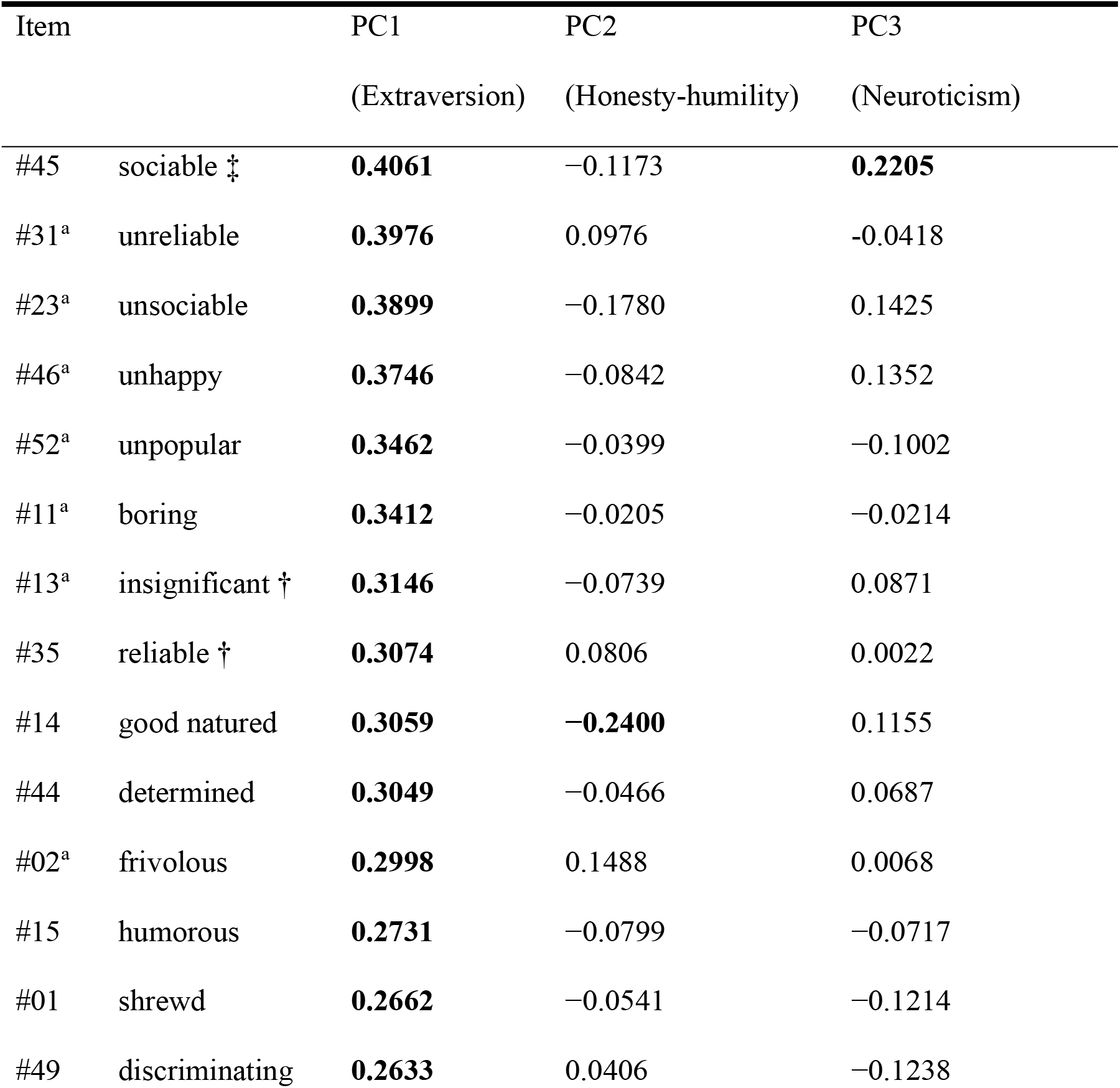
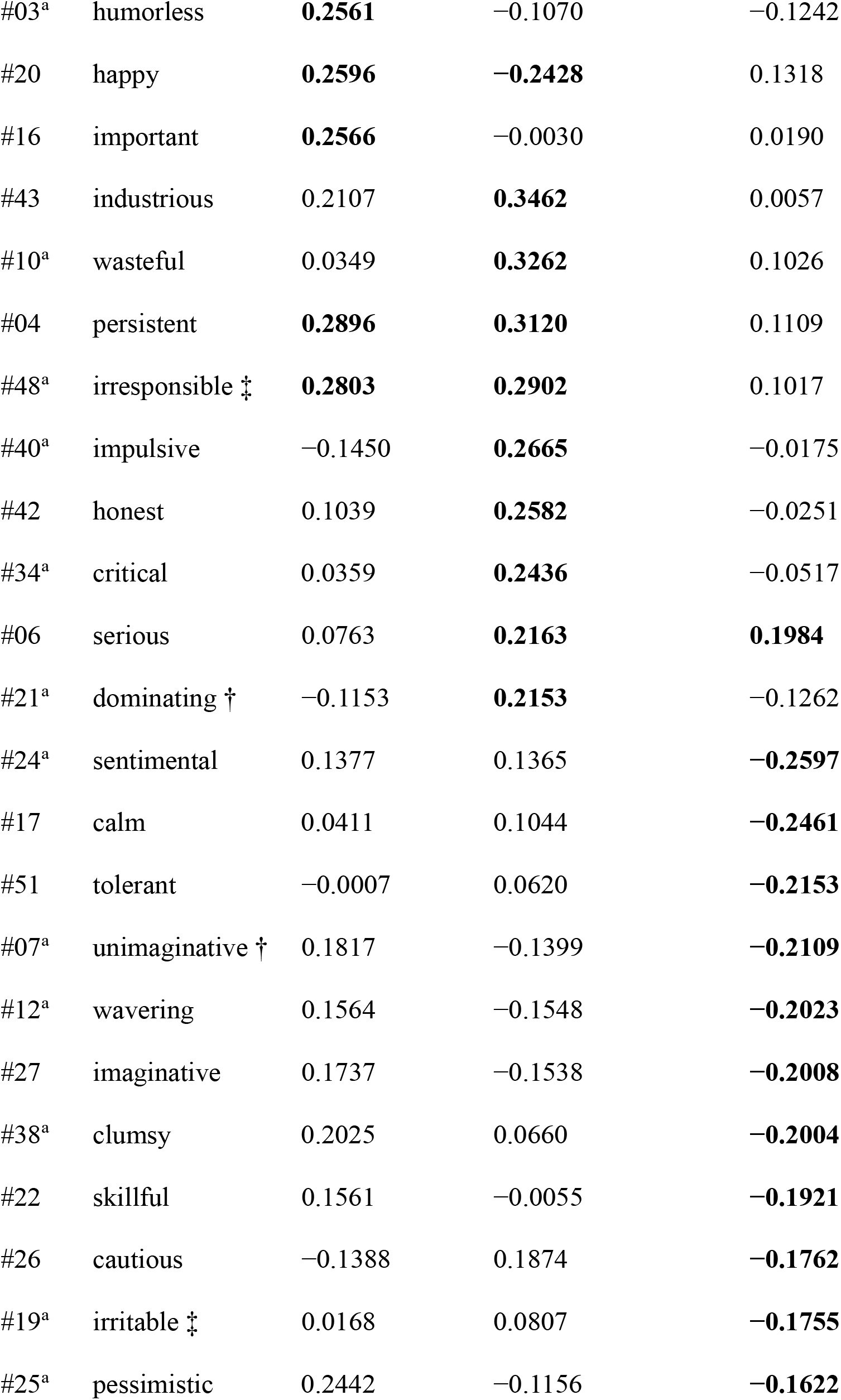
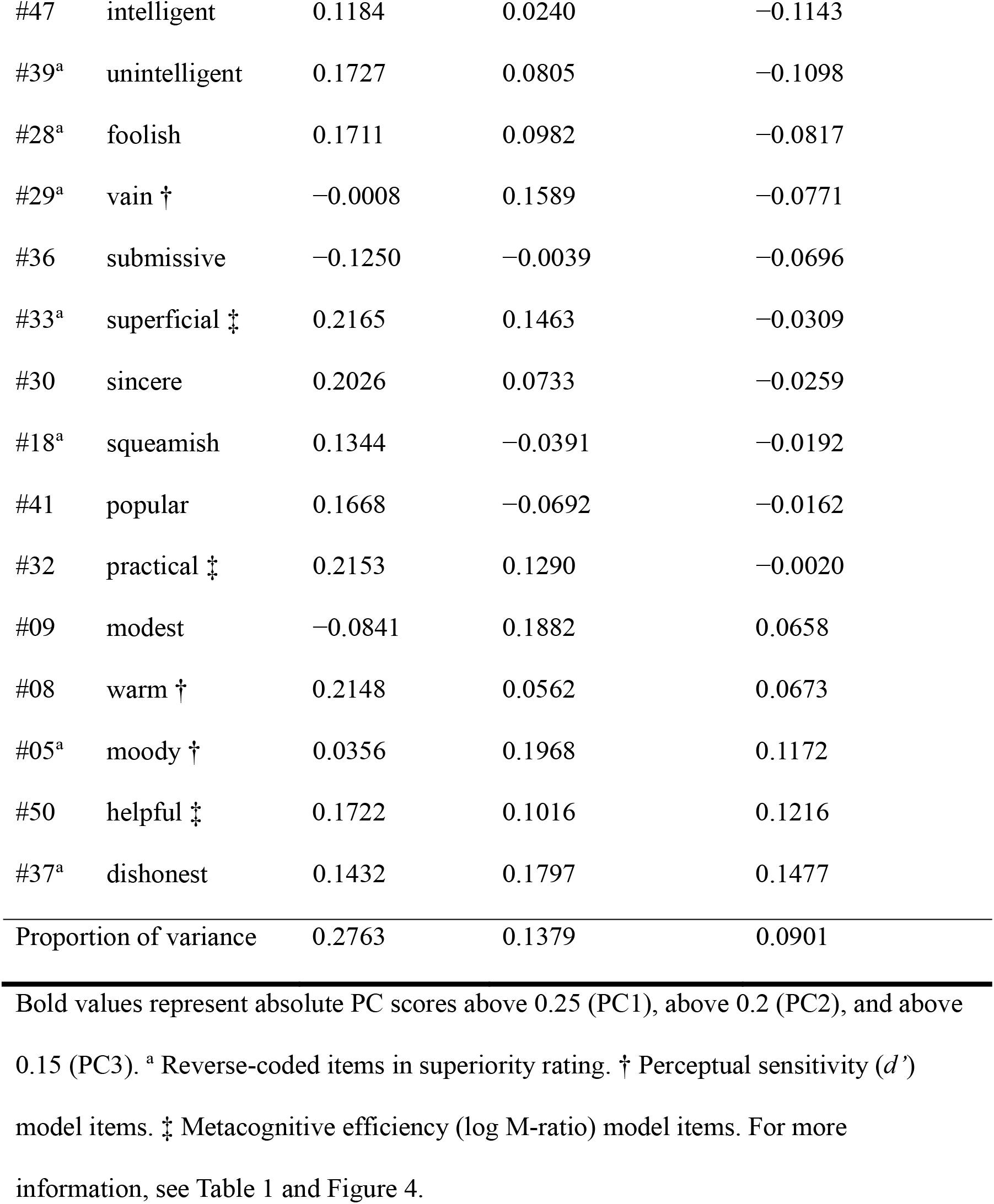
Principal component analysis (PCA) loadings for 52 superiority rating.

| Item |  | PC1 | PC2 | PC3 |
| --- | --- | --- | --- | --- |
|  |  | (Extraversion) | (Honesty-humility) | (Neuroticism) |
| #45 | sociable ‡ | **0.4061** | −0.1173 | **0.2205** |
| #31<sup>a</sup> | unreliable | **0.3976** | 0.0976 | −0.0418 |
| #23<sup>a</sup> | unsociable | **0.3899** | −0.1780 | 0.1425 |
| #46<sup>a</sup> | unhappy | **0.3746** | −0.0842 | 0.1352 |
| #52<sup>a</sup> | unpopular | **0.3462** | −0.0399 | −0.1002 |
| #11<sup>a</sup> | boring | **0.3412** | −0.0205 | −0.0214 |
| #13<sup>a</sup> | insignificant † | **0.3146** | −0.0739 | 0.0871 |
| #35 | reliable † | **0.3074** | 0.0806 | 0.0022 |
| #14 | good natured | **0.3059** | **−0.2400** | 0.1155 |
| #44 | determined | **0.3049** | −0.0466 | 0.0687 |
| #02<sup>a</sup> | frivolous | **0.2998** | 0.1488 | 0.0068 |
| #15 | humorous | **0.2731** | −0.0799 | −0.0717 |
| #01 | shrewd | **0.2662** | −0.0541 | −0.1214 |
| #49 | discriminating | **0.2633** | 0.0406 | −0.1238 |
| #03<sup>a</sup> | humorless | **0.2561** | −0.1070 | −0.1242 |
| #20 | happy | **0.2596** | **−0.2428** | 0.1318 |
| #16 | important | **0.2566** | −0.0030 | 0.0190 |
| #43 | industrious | 0.2107 | **0.3462** | 0.0057 |
| #10<sup>a</sup> | wasteful | 0.0349 | **0.3262** | 0.1026 |
| #04 | persistent | **0.2896** | **0.3120** | 0.1109 |
| #48<sup>a</sup> | irresponsible ‡ | **0.2803** | **0.2902** | 0.1017 |
| #40<sup>a</sup> | impulsive | −0.1450 | **0.2665** | −0.0175 |
| #42 | honest | 0.1039 | **0.2582** | −0.0251 |
| #34<sup>a</sup> | critical | 0.0359 | **0.2436** | −0.0517 |
| #06 | serious | 0.0763 | **0.2163** | **0.1984** |
| #21<sup>a</sup> | dominating † | −0.1153 | **0.2153** | −0.1262 |
| #24<sup>a</sup> | sentimental | 0.1377 | 0.1365 | **−0.2597** |
| #17 | calm | 0.0411 | 0.1044 | **−0.2461** |
| #51 | tolerant | −0.0007 | 0.0620 | **−0.2153** |
| #07<sup>a</sup> | unimaginative † | 0.1817 | −0.1399 | **−0.2109** |
| #12<sup>a</sup> | wavering | 0.1564 | −0.1548 | **−0.2023** |
| #27 | imaginative | 0.1737 | −0.1538 | **−0.2008** |
| #38<sup>a</sup> | clumsy | 0.2025 | 0.0660 | **−0.2004** |
| #22 | skillful | 0.1561 | −0.0055 | **−0.1921** |
| #26 | cautious | −0.1388 | 0.1874 | **−0.1762** |
| #19<sup>a</sup> | irritable ‡ | 0.0168 | 0.0807 | **−0.1755** |
| #25<sup>a</sup> | pessimistic | 0.2442 | −0.1156 | **−0.1622** |
| #47 | intelligent | 0.1184 | 0.0240 | −0.1143 |
| #39<sup>a</sup> | unintelligent | 0.1727 | 0.0805 | −0.1098 |
| #28<sup>a</sup> | foolish | 0.1711 | 0.0982 | −0.0817 |
| #29<sup>a</sup> | vain † | −0.0008 | 0.1589 | −0.0771 |
| #36 | submissive | −0.1250 | −0.0039 | −0.0696 |
| #33<sup>a</sup> | superficial ‡ | 0.2165 | 0.1463 | −0.0309 |
| #30 | sincere | 0.2026 | 0.0733 | −0.0259 |
| #18<sup>a</sup> | squeamish | 0.1344 | −0.0391 | −0.0192 |
| #41 | popular | 0.1668 | −0.0692 | −0.0162 |
| #32 | practical ‡ | 0.2153 | 0.1290 | −0.0020 |
| #09 | modest | −0.0841 | 0.1882 | 0.0658 |
| #08 | warm † | 0.2148 | 0.0562 | 0.0673 |
| #05<sup>a</sup> | moody † | 0.0356 | 0.1968 | 0.1172 |
| #50 | helpful ‡ | 0.1722 | 0.1016 | 0.1216 |
| #37<sup>a</sup> | dishonest | 0.1432 | 0.1797 | 0.1477 |
| Proportion of variance |  | 0.2763 | 0.1379 | 0.0901 |

Figure 5A presents the weighted total feature importance (machine learning feature importance [Table 1] × PCA loadings for the superiority rating items [Table 2]) for each PC and model. The PC1 importance was comparable between the *d’* and log M-ratio model items. However, the *d’* and log M-ratio model items weigh more on PC2 and PC3, respectively. Furthermore, we confirmed that interindividual correlations follow a similar “common yet dissociable” pattern between PCs and SDT parameter estimates (Fig. 5B). We further confirmed the generic (machine learning irrelevant) relationships between the PCs and SDT measures. The correlation between *d’* and superiority rating PCA score was significant in PC1 (*rho* = −0.5545, *p* = 0.0005), while no significant correlations were found in PC2 (*rho* = 0.2603, *p* = 0.1196) and PC3 (*rho* = 0.0462, *p* = 0.7853) (Fig. 5B top row). The correlations between the log M-ratio and superiority rating PCA score were significant in PC1 (*rho* = −0.4388, *p* = 0.0070) and PC3 (*rho* = −0.3812, *p* = 0.0205), while no significant correlation was found in PC2 (*rho* = −0.0363, *p* = 0.8309) (Fig. 5B bottom row).

**Figure 5.**
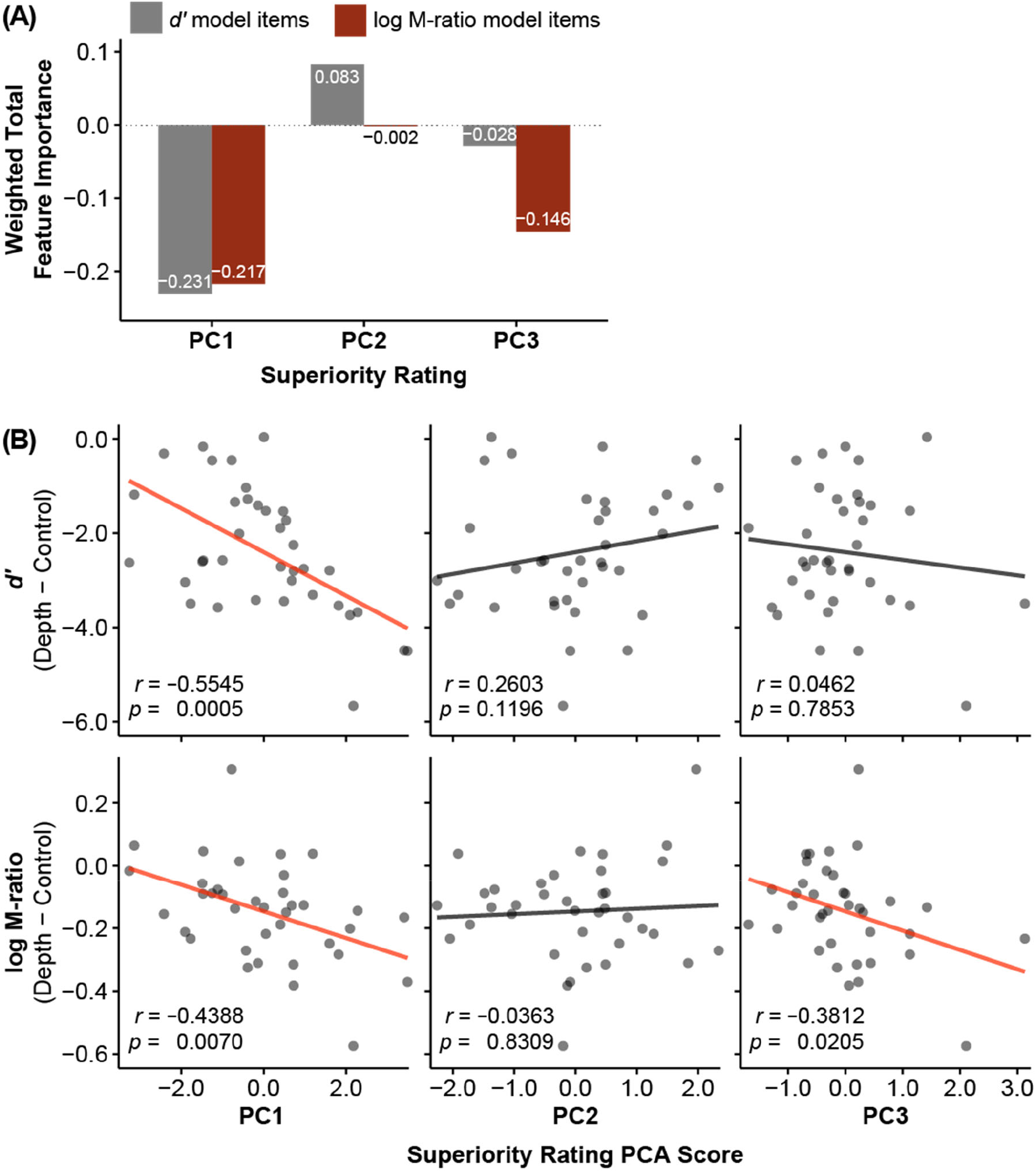
Latent architecture underlying machine-learning-selected superiority rating items. (A) Weighted total feature importance values (machine learning feature importance and PCA loadings products) between the models were comparable in PC1 but dissociable in PC2 and PC3. The results indicate that codes related to SI were overlapping yet dissociable between the visual illusion magnitude (*d’*) and metacognitive performance (log M-ratio). (B) Generic (machine learning irrelevant) relationships between the three PC scores and *d’* (top row) and between the three PC scores and log M-ratio (bottom row). Transparent dots represent individual data points. Transparent lines represent linear regression fit using ordinary least squares.

## Discussion

Leveraging hierarchical Bayesian estimation and machine learning of SDT measures, we aimed to determine how SI influences visual illusion magnitude and metacognitive performance. SI of oneself over an average peer is suggestively crucial for a healthy mental state and behavior (4, 43). However, whether such SI involves low-level perceptual tasks has remained elusive so far. Our behavioral results revealed that SI correlated with visual illusion magnitude and metacognitive ability. Next, our machine learning approach further uncovered the latent architecture behind them. Although the identical superiority feature affected visual illusion magnitude and metacognitive performance, the distinct superiority features simultaneously impacted them. Perception and metacognition are thus liable to influences from both overlapping and separable superiority features. SI might have various psychological benefits (3, 4, 6-10) and exert concurrent biasing effects on visual perception and metacognition, perhaps due to its illusory and self-affirmative belief.

Our findings are in good agreement with recent studies suggesting that global (i.e., general self-belief) and local (i.e., trial-wise decision evaluation) metacognition closely interact, forming a hierarchical structure that impacts mental health (44). They suggested that global self-beliefs bias local confidence, while local confidence helps form global self-beliefs. SI and trial-wise metacognition were closely related, perhaps because the hierarchical structure embeds them as reciprocally-connected layers. SI might accordingly exert a top–down influence on within-hierarchy local metacognition while simultaneously biasing visual illusion strength via a different route, proven by the dissociable contributions of SI features to visual illusion magnitude and local metacognitive performance.

The self-affirmative SI features contributed to perceptual and metacognitive performance. Human variation in subjective superiority in each feature might reflect one’s belief (or priority) of being superior in a given domain (19), which eventually forging individual differences in behavioral heuristics that regulate diverse information processing layers. Human striving to maintain positive self-regard might be a significant source of top-down bias for perceptual capacity to handle contextual information (i.e., the degree to incorporate contexts into visual percepts) and metacognitive ability to monitor self-performance (i.e., the degree of illusory confidence in own perceptual ability).

We identified the *three* features for SI using trait words derived from Rosenberg et al. (45). The authors suggested that the primary components underlying personality impression were *two* (competence and warmth) (for a review, see 46), our results thus appeared to be inconsistent with their dimension number-related findings. However, the *impression of others* and *assessment of own traits* might simply be different things. When people judge social groups, warmth and competence evaluations negatively correlate (47), implying the use of a simplified judgment. Furthermore, Beer and Watson (48) described the convergence tendency of trait dimensions in peer ratings compared to self-ratings. These findings suggest that people use heuristics and judge others based on simplified trait structures. In other words, people might make scrupulous, albeit self-serving, appraisals of their characteristics, resulting in judgments based on elaborated trait structures (49).

Our findings demonstrated shared, yet dissociable, influences of SI on perceptual and metacognitive performance. Extraversion (PC1) is a core feature that affecting *both* visual perception and metacognition, while others do not. Subjective superiority in extraversion was predictive of visual illusion magnitude and metacognitive ability, possibly via lower sensitivity (50-52) and overconfidence (53, 54), respectively. However, lower sensitivity and overconfidence might not be as disparate as it first seems. In fact, they could reflect the two sides of the same coin as in the case of the Dunning–Kruger effect (43, 55), indicating poor performers’ overestimation of their ability (56-58).

Furthermore, honesty-humility (PC2) and neuroticism (PC3) impacted *either* visual illusion strength or metacognitive performance but not both. However, the difference between their contribution to the predictive models was striking. While honesty-humility was predictive of visual illusion magnitude, consistent with the findings showing the correlation between honesty-humility and less dependence on contextual information (59, 60), it contributed to the prediction model relatively weakly (Fig. 5A). Instead, neuroticism contributed to the prediction model more substantially, approximately twice as much as honesty-humility. Therefore, neuroticism might be more operative than honesty-humility in dissociating superiority features and behavioral performance. Although it is well known that neuroticism exhibits fundamental roles in a wide array of health and life outcomes (61), it is surprising that the effect of neuroticism-related metacognition was present even in the perceptual domain.

In conclusion, we discovered that SI correlated with visual illusion strength and metacognitive performance. Moreover, using machine learning, we unveiled their latent architecture predictive of visual perception and metacognition. A significant limitation of our study is that we did not incorporate other classes of visual illusion. How SI influences behavior might hinge on the illusion type (32). In addition, the present findings potentially do not generalize to females as we included only male participants. However, a recent meta-analysis showed that SI per se is constant across gender groups (9). Although further research is warranted to resolve these issues, we suggest that SI is a cardinal cognitive bias that involves a vast assortment of behavior as an illusion is imperative for humans to somehow thrive in a world of ambiguity.

## Materials and Methods

### Ethics statement

The study was approved by the Committee of Ethics, National Institutes for Quantum Science and Technology, Japan. The procedures used in this study adhere to the tenets of the Declaration of Helsinki. All participants provided written informed consent prior to participation.

### Participants

Thirty-seven males participated in this study (mean age: 23.3 ± 3.1 years [1 SD]; range: 20–32 years). All had normal or corrected-to-normal vision and did not report any known neurological or psychiatric conditions.

### Stimuli and procedure

We presented stimuli using E-Prime 2.0 (Psychology Software Tools, PA, USA). Participants viewed stimuli on a 24-inch LCD monitor at a distance of 60 cm. We presented all stimuli on a gray background.

#### Superiority rating task

We successively presented personality trait words on the center of the screen with a visual analog scale (VAS) on the bottom (Fig. 1A). We asked participants to rate the extent to which each personality trait word would describe them by comparing themselves with an (imaginary) average peer using a VAS with a step of 0.05 (score ranges from −1 [much less than the average] through 0 [approximately the same as the average] to 1 [much more than the average]). We used 26 desirable, 26 undesirable, and eight filler words from previous studies (5, 45) in randomized order across the participants. Undesirable-word scores were reverse-coded. Scores beyond zero indicate the subjective superiority of the participants compared to an average person (and vice versa).

#### Ponzo illusion task

We used a black disc (4.6 to 6.7° diameter, randomized across trials) presented at 8.8° to the left and right of the fixation point centered on the screen, as a stimulus to measure the Ponzo illusion (Fig. 1B). The experiment displayed two background image conditions: discs presented on a uniform gray background or a 3D-textured image containing linear-perspective, pictorial depth cues (control and depth cue conditions, respectively).

Each trial comprised the following steps: presentation of a fixation point (500–1000 ms, randomized across trials) followed by a black disc on one side (1000 ms), blank screen (1000 ms), a black disc on the other side (1000 ms), blank screen (300 ms), and two response displays. First, we asked the participants to judge whether the two discs were of the same size by pressing a corresponding response pad. Second, the participants had to rate their confidence for the first decision by pressing a corresponding key on a scale of 1 (very unconfident) to 4 (very confident). The participants carried out 320 trials, consisting of 128, 128, 32, and 32 trials where the “distant” disc was equal to, 20% smaller, 5% smaller, and 5% larger in diameter than the other, respectively. The 5% larger/smaller sets represented filler trials and were not analyzed further. Half of 320 trials were performed under depth cue, and the other half under control conditions. In the case of the depth cue conditions, the left wall was apparently “close” on half of the trials, and the right wall was apparently “close” on the other half. We always presented the first disc on the “close” side of the wall. Due to the uniform background, no markedly “distant” or “close” disc could be distinguished under the control (but not the depth cue) conditions. The trial order was pseudo-randomized across the trials such that all conditions appeared in every 40 trials. The participants took a few-minute break after performing 160 trials.

### Hierarchical Bayesian estimation of SDT measures

To estimate metacognitive efficiency, we computed log(meta-*d’*/*d’*), where *d’* is an SDT measure of type 1 first-order sensitivity (i.e., *perceptual* sensitivity) and meta-*d’* is a measure of type 2 *metacognitive* sensitivity (1), representing a measure of the ability to distinguish between correct and incorrect judgments. Meta-*d’*/*d’*, also called M-ratio, is a measure of metacognitive *efficiency*, compensating for the intrinsic correlation between meta-*d’* and *d’*. Meta-*d’* equals to *d’* (i.e., M-ratio = 1 and log M-ratio = 0) represents that the observer is metacognitively “optimal,” using all the available information for the type 1 task to the type 2 task. However, people are typically not fully aware of the accuracy of a decision; observers often display metacognitive *inefficiency* (i.e., M-ratio < 1 and log M-ratio < 0) (62). In contrast, observers occasionally exhibit *superefficiency* (i.e., M-ratio > 1 and log M-ratio > 0) in that they seemingly use more information than the theoretical maximum (63, 64). Although superefficiency is not well understood, the nonoptimal metacognition (i.e., either inefficiency or superefficiency) implies (at least partially) distinct mechanisms for first-order decisions and confidence ratings.

We performed hierarchical Bayesian estimation of the SDT parameters using Markov chain Monte Carlo sampling (3 chains of 10,000 samples, and 1,000 burn-in samples) to incorporate within- and between-subject uncertainty and avoid edge-correction confounds (41). The hierarchical Bayesian approach allows for recovering accurate metacognitive efficiency estimates from confidence ratings even at low *d’* values, where commonly used alternatives fail. This characteristic benefits our Ponzo illusion task with an inherently low perceptual sensitivity (i.e., illusion leads to poor discrimination performance). We performed statistical analyses on log M-ratio (instead of M-ratio) to ensure that a unit of distance along an axis represents an equal weight relative to the optimal value of meta-*d’*/*d’* = 1 (41, 65).

### Machine learning using a relaxed elastic net

We created a prediction model using a machine learning technique to examine which set of superiority rating items explains best each SDT parameter estimate of the Ponzo illusion. We performed a relaxed elastic net, a two-step elastic net regression similar to a relaxed Lasso (66). Relaxed elastic net regression creates a regularized regression model by performing variable (superiority rating item) selection using the standard elastic net (67), then determines weight coefficients for the selected variables using ridge regression. This procedure attenuates overfitting and multicollinearity by shrinking variance and results in more reliable estimates than conventional linear regression using ordinary least squares. We created two models: one to predict *d’* and another to predict log M-ratio from 52-item superiority ratings. All variables included in the models were standardized to have zero mean and one variance. We performed a relaxed elastic net regression with LOOCV: we used α ∈ [0.1, 1.0] (a hyperparameter controlling the trade-off between the L1 and L2 penalties) with a step of 0.1 and λ ∈ 10^[-3,3]^ (a regularization hyperparameter) with a step of 2/33 in the initial elastic net, then zero α and the best-tuned λ (from the initial elastic net) to optimize the weight vector of the selected items in the following ridge regression. This two-step procedure effectively reduces the dimensionality of the superiority rating items related to the visual illusion SDT parameter estimates through variable selection while providing more optimal weight estimates than standard elastic net regression (see 68).

### PCA

We performed a PCA with singular value decomposition on 52-item superiority ratings to estimate latent components underlying SI. We performed a parallel analysis using unweighted least squares to find an optimal number of PCs (42). Next, to examine the relationship between machine-learning-selected superiority rating items and SDT parameter estimates, we calculated an index called weighted total feature importance, representing the relative contribution of each PC to each machine learning model by taking the matrix product of machine learning feature importance and PCA loadings. Higher (absolute) values indicate a higher contribution of that particular PC to the prediction model. Moreover, we examined correlations between PC scores and SDT parameter estimates to confirm the generic relationship between superiority rating PCs and SDT parameter estimates.

### Statistical inference

We set the statistical thresholds at α = 0.05 for superiority ratings and type 1 SDT measures (*d’* and *C*) and at 95% highest density interval (HDI) of posterior distributions for group-level hierarchical Bayesian type 2 SDT parameter estimates (M-ratio and log M-ratio). To accurately capture the effects of the visual illusion, we calculated between-condition differences for the SDT parameter estimates (depth condition − control condition). A negative difference value indicated a higher Ponzo illusion magnitude (*d’*), a more liberal criterion under the illusion (*C*), or lower illusion-induced metacognitive performance (M-ratio and log M-ratio). We used parameter estimates from single-subject Bayesian model fits for correlation and individual difference analyses. We assessed correlations using Spearman’s rho and set the significance threshold at α = 0.05.

## Acknowledgments

We thank Keita Yokokawa and Keisuke Takahata for their assistance in preparing Ponzo illusion task. This study was supported by grants from JST Moonshot R&D Grant (JPMJMS2295-01 to M.Y.), and the JSPS KAKENHI Grant (19H04433 to D.M.; 20H05711, 22H01108, and 22K18265 to M.Y.).

## Competing interests

None of the authors have any potential conflicts of interest.

## Data availability statements

Data and code required to reproduce the results in this paper are found at https://github.com/dicemt/*** (upon publication)

## Notes

### Competing Interest Statement

The authors have declared no competing interest.

## References

1. Maniscalco B, Lau H (2012) A signal detection theoretic approach for estimating metacognitive sensitivity from confidence ratings. Conscious Cogn 21(1):422–430.

2. Shekhar M, Rahnev D (2021) The nature of metacognitive inefficiency in perceptual decision making. Psychol Rev 128(1):45–70.

3. Taylor SE, Brown JD (1994) Positive illusions and well-being revisited: Separating fact from fiction. Psychol Bull 116(1):21–27.

4. Taylor SE, Brown JD (1988) Illusion and well-being: A social psychological perspective on mental health. Psychol Bull 103(2):193–210.

5. Yamada M, et al. (2013) Superiority illusion arises from resting-state brain networks modulated by dopamine. Proc Natl Acad Sci USA 110(11):4363–4367.

6. Taylor SE, Kemeny ME, Reed GM, Bower JE, Gruenewald TL (2000) Psychological resources, positive illusions, and health. Am Psychol 55(1):99–109.

7. Gana K, Alaphilippe D, Bailly N (2004) Positive illusions and mental and physical health in later life. Aging Ment Health 8(1):58–64.

8. Kurman J (2003) Why is self-enhancement low in certain collectivist cultures?: An investigation of two competing explanations. J Cross-Cult Psychol 34(5):496–510.

9. Zell E, Strickhouser JE, Sedikides C, Alicke MD (2020) The better-than-average effect in comparative self-evaluation: A comprehensive review and meta-analysis. Psychol Bull 146(2):118–149.

10. Goetz T, Ehret C, Jullien S, Hall NC (2006) Is the grass always greener on the other side? Social comparisons of subjective well-being. J Posit Psychol 1(4):173–186.

11. Sedikides C, Horton RS, Gregg AP (2007) The why’s the limit: Curtailing self-enhancement with explanatory introspection. J Pers 75(4):783–824.

12. Colvin CR, Block J, Funder DC (1995) Overly positive self-evaluations and personality: Negative implications for mental health. J Pers Soc Psychol 68(6):1152–1162.

13. Bortolotti L, Antrobus M (2015) Costs and benefits of realism and optimism. Curr Opin Psychiatry 28(2).

14. Light N, Fernbach PM, Rabb N, Geana MV, Sloman SA (2022) Knowledge overconfidence is associated with anti-consensus views on controversial scientific issues. Sci Adv 8(29):eabo0038.

15. Hoorens V (1993) Self-enhancement and superiority biases in social comparison. Eur Rev Soc Psychol 4(1):113–139.

16. Svenson O (1981) Are we all less risky and more skillful than our fellow drivers? Acta Psychol (Amst) 47(2):143–148.

17. Tappin BM, McKay RT (2016) The illusion of moral superiority. Soc Psychol Pers Sci 8(6):623–631.

18. Heine SJ, Lehman DR (1997) The cultural construction of self-enhancement: An examination of group-serving biases. J Pers Soc Psychol 72(6):1268–1283.

19. Sedikides C, Gaertner L, Toguchi Y (2003) Pancultural self-enhancement. J Pers Soc Psychol 84(1):60–79.

20. Wu S (2018) No Lake Wobegon in Beijing? The impact of culture on the perception of relative ranking. Appl Cogn Psychol 32(2):192–199.

21. Lee DY, Park SH, Uhlemann MR (2002) Self and other ratings of Canadian and Korean groups of mental health professionals and their clients. Psychol Rep 90(2):667–676.

22. Witkin HA, Moore CA, Goodenough DR, Cox PW (1977) Field-dependent and field-independent cognitive styles and their educational implications. Rev Edu Res 47(1):1–64.

23. Witkin HA, Goodenough DR (1977) Field dependence and interpersonal behavior. Psychol Bull 84(4):661–689.

24. Kitayama S, Duffy S, Kawamura T, Larsen JT (2003) Perceiving an object and its context in different cultures: A cultural look at new look. Psychol Sci 14(3):201–206.

25. MacLeod CM, Jackson RA, Palmer J (1986) On the relation between spatial ability and field dependence. Intelligence 10(2):141–151.

26. Shoshina II, Shelepin YE (2014) Effectiveness of discrimination of the sizes of line segments by humans with different cognitive style parameters. Neurosci Behav Physiol 44(7):748–753.

27. Zhang L-f (2004) Field-dependence/independence: cognitive style or perceptual ability?––validating against thinking styles and academic achievement. Pers Individ Dif 37(6):1295–1311.

28. Sih A, Del Giudice M (2012) Linking behavioural syndromes and cognition: a behavioural ecology perspective. Philos Trans R Soc B Biol Sci 367(1603):2762–2772.

29. Kozhevnikov M (2007) Cognitive styles in the context of modern psychology: Toward an integrated framework of cognitive style. Psychol Bull 133(3):464–481.

30. Cuneo F, Antonietti J-P, Mohr C (2018) Unkept promises of cognitive styles: A new look at old measurements. PLoS One 13(8):e0203115.

31. Stark E, Stacey J, Mandy W, Kringelbach ML, Happé F (2021) Autistic cognition: Charting routes to anxiety. Trends Cogn Sci 25(7):571–581.

32. Gregory RL (1997) Visual illusions classified. Trends Cogn Sci 1(5):190–194.

33. Eagleman DM (2001) Visual illusions and neurobiology. Nat Rev Neurosci 2(12):920–926.

34. Yildiz GY, Sperandio I, Kettle C, Chouinard PA (2022) A review on various explanations of Ponzo-like illusions. Psychon Bull Rev 29(2):293–320.

35. Murray SO, Boyaci H, Kersten D (2006) The representation of perceived angular size in human primary visual cortex. Nat Neurosci 9(3):429–434.

36. Fang F, Boyaci H, Kersten D, Murray SO (2008) Attention-dependent representation of a size illusion in human V1. Curr Biol 18(21):1707–1712.

37. Bosking WH, Zhang Y, Schofield B, Fitzpatrick D (1997) Orientation selectivity and the arrangement of horizontal connections in tree shrew striate cortex. J Neurosci 17(6):2112–2127.

38. Gilbert C, Wiesel T (1989) Columnar specificity of intrinsic horizontal and corticocortical connections in cat visual cortex. J Neurosci 9(7):2432–2442.

39. Schwarzkopf DS, Song C, Rees G (2011) The surface area of human V1 predicts the subjective experience of object size. Nat Neurosci 14(1):28–30.

40. Barrett AB, Dienes Z, Seth AK (2013) Measures of metacognition on signal-detection theoretic models. Psychol Methods 18(4):535–552.

41. Fleming SM (2017) HMeta-d: hierarchical Bayesian estimation of metacognitive efficiency from confidence ratings. Neurosci Conscious 2017(1).

42. Horn JL (1965) A rationale and test for the number of factors in factor analysis. Psychometrika 30(2):179–185.

43. Kruger J, Dunning D (1999) Unskilled and unaware of it: How difficulties in recognizing one’s own incompetence lead to inflated self-assessments. J Pers Soc Psychol 77(6):1121–1134.

44. Seow TXF, Rouault M, Gillan CM, Fleming SM (2021) How local and global metacognition shape mental health. Biol Psychiatry 90(7):436–446.

45. Rosenberg S, Nelson C, Vivekananthan PS (1968) A multidimensional approach to the structure of personality impressions. J Pers Soc Psychol 9(4):283–294.

46. Fiske ST, Cuddy AJC, Glick P (2007) Universal dimensions of social cognition: warmth and competence. Trends Cogn Sci 11(2):77–83.

47. Fiske ST, Cuddy AJC, Glick P, Xu J (2002) A model of (often mixed) stereotype content: Competence and warmth respectively follow from perceived status and competition. J Pers Soc Psychol 82(6):878–902.

48. Beer A, Watson D (2008) Personality judgment at zero acquaintance: Agreement, assumed similarity, and implicit simplicity. J Pers Assess 90(3):250–260.

49. Klein SB, Kihlstrom JF (1986) Elaboration, organization, and the self-reference effect in memory. J Exp Psychol Gen 115(1):26–38.

50. Blumenthal TD (2001) Extraversion, attention, and startle response reactivity. Pers Individ Dif 31(4):495–503.

51. Fine BJ, Kobrick JL (1976) Note on the relationship between introversion-extraversion, field-dependence-independence and accuracy of visual target detection. Percept Mot Skills 42(3, Pt 1):763-766.

52. Harkins S, Geen RG (1975) Discriminability and criterion differences between extraverts and introverts during vigilance. J Res Pers 9(4):335–340.

53. Schaefer PS, Williams CC, Goodie AS, Campbell WK (2004) Overconfidence and the Big Five. J Res Pers 38(5):473–480.

54. Vaughan-Johnston TI, MacGregor KE, Fabrigar LR, Evraire LE, Wasylkiw L (2021) Extraversion as a moderator of the efficacy of self-esteem maintenance strategies. Pers Soc Psychol Bull 47(1):131–145.

55. Kruger J, Dunning D (2002) Unskilled and unaware--but why? A reply to Krueger and Mueller (2002). J Pers Soc Psychol 82(2):189–192.

56. Gignac GE, Zajenkowski M (2020) The Dunning-Kruger effect is (mostly) a statistical artefact: Valid approaches to testing the hypothesis with individual differences data. Intelligence 80(101449.

57. McIntosh RD, Fowler EA, Lyu T, Della Sala S (2019) Wise up: Clarifying the role of metacognition in the Dunning-Kruger effect. J Exp Psychol Gen 148(11):1882–1897.

58. Burson KA, Larrick RP, Klayman J (2006) Skilled or unskilled, but still unaware of it: How perceptions of difficulty drive miscalibration in relative comparisons. J Pers Soc Psychol 90(1):60–77.

59. Wendler K, Liu J, Zettler I (2018) Honesty-humility interacts with context perception in predicting task performance and organizational citizenship behavior. J Pers Psychol 17(4):161–171.

60. Wiltshire J, Bourdage JS, Lee K (2014) Honesty-humility and perceptions of organizational politics in predicting workplace outcomes. J Bus Psychol 29(2):235–251.

61. Widiger TA, Oltmanns JR (2017) Neuroticism is a fundamental domain of personality with enormous public health implications. World Psychiatry 16(2):144–145.

62. Shekhar M, Rahnev D (2021) Sources of metacognitive inefficiency. Trends Cogn Sci 25(1):12–23.

63. Rahnev D (2021) Visual metacognition: Measures, models, and neural correlates. Am Psychol 76(9):1445–1453.

64. Fleming SM, Daw ND (2017) Self-evaluation of decision-making: A general Bayesian framework for metacognitive computation. Psychol Rev 124(1):91–114.

65. Morales J, Lau H, Fleming SM (2018) Domain-general and domain-specific patterns of activity supporting metacognition in human prefrontal cortex. J Neurosci 38(14):3534–3546.

66. Meinshausen N (2007) Relaxed Lasso. Comput Stats Data Anal 52(1):374–393.

67. Zou H, Hastie T (2005) Regularization and variable selection via the elastic net. J R Stat Soc Series B Stat Methodol 67(2):301–320.

68. Kobak D, et al. (2021) Sparse reduced-rank regression for exploratory visualisation of paired multivariate data. J R Stat Soc Ser C Appl Stat 70(4):980–1000.

